# High affinity cross-context cellular assays reveal novel protein-protein interactions of peripheral myelin protein of 22 kDa

**DOI:** 10.1101/2025.09.03.673966

**Authors:** D Stausberg, S Moshkovskii, FA Arlt, R Fledrich, RM Stassart, KA Nave, H Urlaub, D Ewers, MW Sereda

## Abstract

Peripheral Myelin Protein 22 (PMP22) is a tetraspan membrane protein whose altered dosage causes the most common hereditary neuropathy, Charcot-Marie-Tooth disease type 1A (CMT1A). Despite its clinical significance, the physiological functions of PMP22 and the mechanism behind its tightly controlled gene dosage sensitivity remain unknown since over 30 years, in part due to limited knowledge of its protein-protein interactions (PPIs). In fact, integral membrane proteins such as PMP22 are significantly underrepresented in known cellular interactomes, likely due to limited suitability or technical challenges specific to these hydrophobic molecules in the major PPI discovery approaches. Here, we applied a rigorously optimized co-immunoprecipitation and mass spectrometry workflow using the mild detergent DDM and the high affinity ALFA-tag/anti-ALFA nanobody interaction to identify cellular PMP22-associated proteins. In a cross-context approach, we ran our standardized pipeline across multiple cell types including HEK293T, MDCKII epithelial cells, the Schwann cell line MSC80, and primary rat Schwann cells. We confirm known interactors, and uncover distinct, cell type-specific enrichment patterns following functional annotation analysis. Adhesion-related PPIs dominated in MDCKII cells (e.g., CD47, CLDN1, ATP1B1), while in Schwann cells myelin-associated PPIs were enriched. Importantly, we identified novel PPI candidates that may be highly relevant for PMP22 function including enzymes of the *de novo* sphingolipid biosynthesis pathway.

## Introduction

Peripheral Myelin Protein 22 kDa (PMP22) is a small tetraspan transmembrane protein that is predominantly expressed by myelinating Schwann cells (Kitamura *et al*, 1976; Snipes *et al*, 1992; Zanotti *et al*, 2022). *PMP22* has received widespread attention since it was revealed as the disease-causing gene in Charcot-Marie-Tooth disease type 1A (CMT1A), which results from an intrachromosomal gene duplication leading to approximately 1.5-fold overexpression at both the mRNA and protein level (Lupski *et al*, 1991; Raeymaekers *et al*, 1991; Lupski *et al*, 1992; Patel *et al*, 1992; Matsunami *et al*, 1992; Timmerman *et al*, 1992; Valentijn *et al*, 1992; Sereda *et al*, 1996; Li *et al*, 2013; Hertzog & Jacob, 2023). CMT1A, the most prevalent hereditary neuropathy, is characterized by abnormal myelin growth and subsequent axonal loss (Pareyson & Marchesi, 2009; Pisciotta & Shy, 2023). We and others have established abnormal trafficking of the disease protein (Marinko *et al*, 2020) and insufficient differentiation of Schwann cells with dysregulation of (myelin) growth signaling (Fledrich *et al*, 2014, 2019; Fornasari *et al*, 2018; Krauter *et al*, 2024) and lipid metabolism (Vigo *et al*, 2005; Fledrich *et al*, 2012, 2018; Visigalli *et al*, 2020; Zhou *et al*, 2020; Michailidou *et al*, 2023; Capodivento *et al*, 2024; Prior *et al*, 2024), but the normal function of PMP22 and the details of the CMT1A disease mechanism remain largely unclear. Multiple preclinical therapeutic approaches aimed to reduce *PMP22* mRNA levels (Sereda *et al*, 2003; Meyer zu Hörste *et al*, 2007; Chumakov *et al*, 2014; Zhao *et al*, 2018; Prukop *et al*, 2019, 2020; Serfecz *et al*, 2019; Boutary *et al*, 2021, 2025; Gautier *et al*, 2021; Van Lent *et al*, 2023; Yoshioka *et al*, 2023; Espallergues *et al*, 2025). However, the degree of downregulation remains a challenge as the normal dosage window is very narrow, because too little PMP22 (haploinsufficiency) causes hereditary neuropathy with liability to pressure palsies (HNPP) (Adlkofer *et al*, 1995, 1997).

The activation of cellular stress cascades upon misfolding and aggregation of overexpressed PMP22 has been proposed as the primary CMT1A disease mechanism (Notterpek et al., 1999). However, the absence of aggregates and mistrafficking in peripheral nerves of young CMT1A patients (Nishimura *et al*, 1996; Hanemann *et al*, 2000) and suitable CMT1A models (Jouaud *et al*, 2019; Niemann *et al*, 2000) challenge this notion. We note that the PI3K-Akt-mTOR signaling cascade, which is important for myelin growth, is dysregulated in opposite directions in CMT1A and HNPP (Fledrich *et al*, 2014; Krauter *et al*, 2024), although, in the latter, no pathological aggregation should be expected. Thus, gain of a normal function of PMP22 in regulation of myelin biosynthesis seems more likely responsible for the lack of Schwann cell differentiation and subsequent dysmyelination in CMT1A. In line with this scenario, before its role in peripheral myelination was recognized, *PMP22* was discovered as a growth arrest specific gene (*gas3*) in fibroblasts (Schneider *et al*, 1988), and its expression in multiple cell types (Suter *et al*, 1994) and reported roles in progression of various cancers (Winslow *et al*, 2013; Liu & Chen, 2015; Qu *et al*, 2015) suggest a growth-regulatory function of PMP22 in non-myelinating cells.

Protein-protein interactions (PPIs) are essential for the cellular function of most proteins (Stumpf *et al*, 2008; Venkatesan *et al*, 2009; Michaelis *et al*, 2023). PMP22 may thus exert its function(s) both in Schwann cells and other cell types through physical interaction with other proteins, making comprehensive discovery of PMP22’s PPIs an important prerequisite for studies aiming to further elucidate its role in health and disease. Several putative PPIs of endogenous PMP22 have already been detected in peripheral nerves using co-immunoprecipitation (Co-IP) (Dickson *et al*, 2002; Amici *et al*, 2006; Guo *et al*, 2014; Hu *et al*, 2016; Vanoye *et al*, 2019; Zhou *et al*, 2019; D’Urso *et al*, 1999), a powerful method to identify direct or indirect interactions. However, the success of Co-IP largely depends on the availability of specific antibodies and, in the case of membrane proteins such as PMP22, on gentle and efficient solubilization from the lipid environment, and these inherent technical challenges may explain previous conflicting results (Amici *et al*, 2006; Poitelon *et al*, 2018). In cultured cells that can be genetically manipulated easily, the specificity problem can be avoided by fusing the protein of interest to epitope- or affinity-tags that can be used to perform Co-IP or affinity purification (AP) under more stringent conditions. Using Co-IP/AP, previously found interactors such as Myelin protein zero (MPZ) (D’Urso *et al*, 1999) and Calnexin (Canx) (Dickson *et al*, 2002) were confirmed (Fontanini *et al*, 2005; Hara *et al*, 2014; Marinko *et al*, 2021; Pashkova *et al*, 2024), and more PPI candidates identified also in non-myelinating cells (Wilson *et al*, 2002; Guo *et al*, 2014; Hara *et al*, 2014; Hu *et al*, 2016; Zhou *et al*, 2019; Marinko *et al*, 2021). Several efforts using Yeast Two-Hybrid (Y2H) screening or Co-IP-mass spectrometry (MS) to reveal PPIs of single proteins or whole interactomes (Wang *et al*, 2011, 2023; Dittmer *et al*, 2014; Rolland *et al*, 2014; Sahni *et al*, 2015; Luck *et al*, 2020; Haenig *et al*, 2020) or specifically PMP22 interactors (Marinko *et al*, 2021) detected further candidate PPIs of PMP22. However, the limited overlap between the previous results indicates that the identified PPIs are far from covering all of PMP22’s interaction partners, and few PPIs have been found in Schwann cells so far. To establish a solid foundation for functional studies on the role of PMP22 in both Schwann cells and non-myelinating cells, we used an optimized Co-IP-MS approach that employs a novel epitope tag with high affinity to an engineered nanobody, to generate a comprehensive list of PMP22 interaction candidates in various cell types, including Schwann cells.

## Results

To investigate PPIs of PMP22, we used Co-IP-MS from HEK293T (HEK) cells, a strategy that has also been used by Sanders and colleagues (Marinko *et al*, 2021). This easy to transfect cell line that is likely of adrenal origin (Russell *et al*, 1977; Shaw *et al*, 2002; Lin *et al*, 2014) lacks a known functional role of endogenous PMP22. The cell types in which such role has been described show pronounced front-rear (fibroblasts) or apical-basal (epithelial cells, Schwann cells) polarity, while HEK cells remain non-polarized. Thus, HEK cells seem suitable as a model system to investigate basic PPI of PMP22 that are not required for its specialized function in polarized cells. To enable efficient capture of PMP22 in Co-IP, we decided to use the ALFA-tag, which was recently developed by rational design and reported to enable low-background IP by means of the highly specific and high-affinity interaction with anti-ALFA nanobody (Götzke *et al*, 2019). We employed a plasmid that allows expression of C-terminally ALFA-tagged human PMP22 (Fig. 1A) together with a blue fluorescent protein (mTAGBFP2) (Subach *et al*, 2011) driven by a separate promotor to allow for identification of transfected cells via fluorescent microscopy. Following transfection of HEK cells, Western Blot analysis showed ALFA-tag positive bands in the cell lysate that were mostly PNGase F sensitive and Endo H resistant, indicating complex glycosylation after Golgi processing (Fig. 1B). In line with this observation, immunostaining revealed PMP22-ALFA at or near the cell surface (Fig. 1C). Co-IP of membrane proteins requires their efficient extraction from the lipid bilayer with subsequent ultracentrifugation to enable separation from non-interacting, colocalizing proteins. Fig. 1D shows successful solubilization of PMP22-ALFA in all glycosylation states by n-Dodecyl-β-D-maltopyranoside (DDM), a gentle, non-ionic detergent. DDM is known for its superior efficiency in membrane extraction for structural studies (Newstead *et al*, 2008; Stetsenko & Guskov, 2017; Choy *et al*, 2021; Harrison *et al*, 2023; Vénien-Bryan & Fernandes, 2023), and was recently applied in combination with cholesteryl hemisuccinate for successful Co-IP of PMP22 and MPZ (Pashkova *et al*, 2024). Interestingly, that study also reported failure of Co-IP with NP-40 and Triton X-100, two detergents previously used for Co-IP of PMP22 with MPZ and other proteins (Fontanini *et al*, 2005; D’Urso *et al*, 1999). A recent Co-IP-MS study used CHAPS (Marinko *et al*, 2021), a zwitterionic detergent that has been reported to be more efficient in breaking PPIs than NP-40 and Triton X-100 (Labeta *et al*, 1988). To assess its suitability for solubilization of PMP22-ALFA from HEK cell membranes, we tested CHAPS at a concentration of 0.3%, which was used by Sanders and colleagues, as well as at 1%, well above the critical micelle concentration (Chattopadhyay & Harikumar, 1996). Both conditions resulted in extraction of only a small fraction of PMP22-ALFA, specifically of lower molecular weight, while CHAPS failed to solubilize higher molecular weight PMP22-ALFA (Fig. 1D). Unlike DDM, CHAPS therefore seemed not well suited for extracting PMP22-ALFA from the Golgi and the plasma membrane for IP. We thus performed IP using DDM for membrane solubilization and magnetic beads coupled to anti-ALFA nanobody. As inputs we used the supernatant after ultracentrifugation of solubilized HEK cells transiently expressing either PMP22-ALFA or untagged PMP22 as a control. Transfection efficiencies were similar as judged by comparable fluorescence of co-expressed mTAGBFP2 (suppl. Fig. 1). SDS-PAGE with subsequent silver staining showed protein bands resembling PMP22-ALFA in different glycosylation states as previously identified (Fig. 1B) in the eluate of PMP22-ALFA IP, but not of PMP22 IP (Fig. 1E). In line with this observation, label-free quantification (LFQ) MS after chymotrypsin-cleavage detected similar abundance of PMP22 in both inputs and showed enrichment of PMP22 in the PMP22-ALFA IP eluate, while PMP22 was absent from the IP eluate of untagged PMP22 control (Fig. 1F). PMP22 was previously reported to form homo-oligomers (Tobler *et al*, 1999; Mobley *et al*, 2007). To test the Co-IP approach for successful enrichment of a known, probably directly interacting protein, we thus performed PMP22-ALFA IP following co-transfection of HEK cells with a C-terminal fusion of PMP22 with a monomeric GFP. Indeed, we detected PMP22-GFP via Western Blot analysis in the PMP22-ALFA eluate but not in the negative control, demonstrating self-interaction of PMP22 in HEK cell membranes (Fig. 1G).

**Figure 1.**
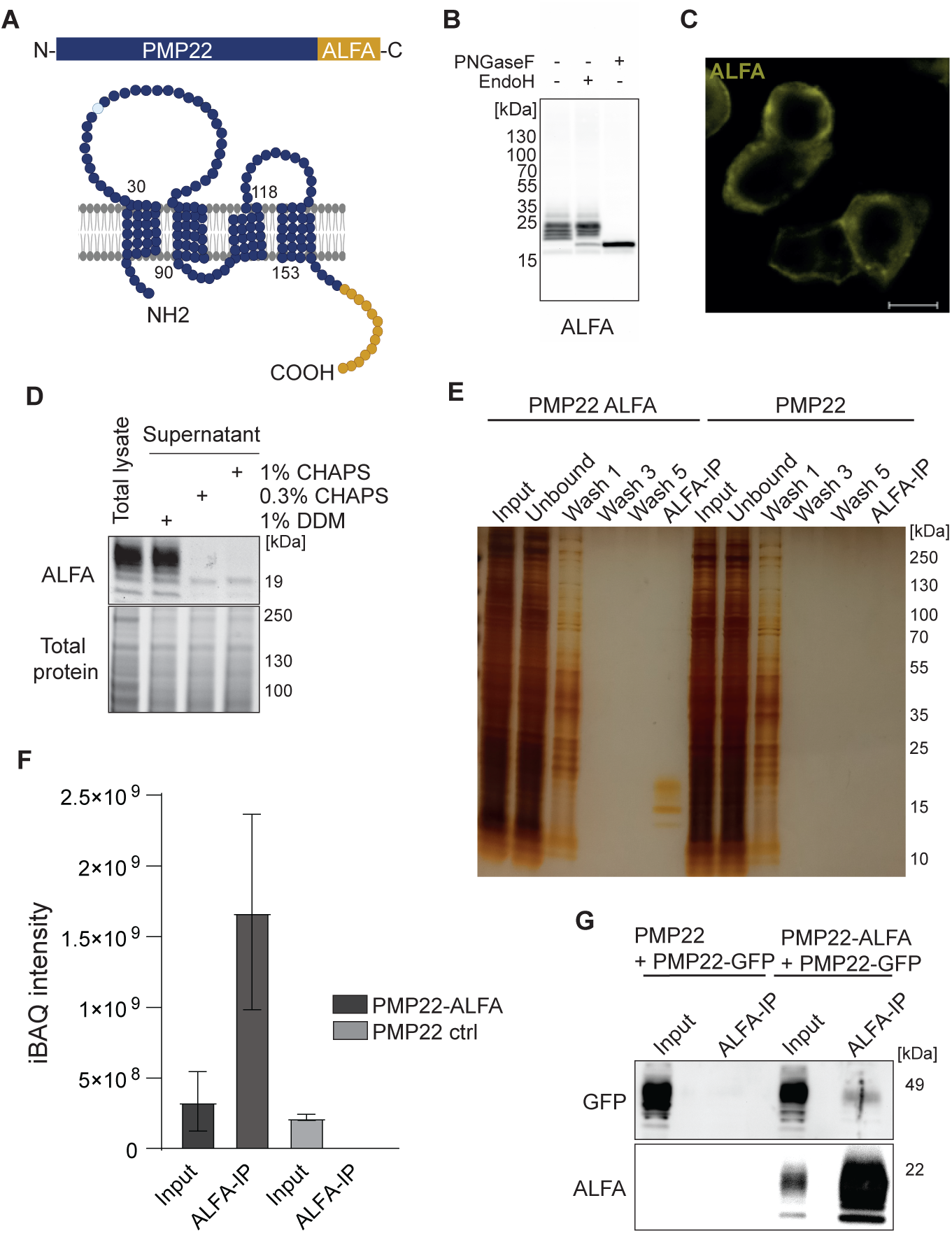
Low-recovery, low-background Co-IP of PMP22 via the ALFA tag. (A) Schematic representation of the PMP22 secondary protein structure with the C-terminal ALFA tag. N-linked glycosylation site at asparagine 41 is highlighted in light blue. (B) Western Blot of HEK cell lysate after PMP22-ALFA transfection subjected to PNGase F and Endo H digestion, showing complex glycosylation of PMP22-ALFA with multiple bands ranging from 19 to 25 kDa. (C) Immunofluorescent staining shows PMP22-ALFA (orange) localized at the plasma membrane in HEK cells. Scale bar is 10 µm. (D) Western Blot analysis of detergent-solubilized PMP22-ALFA comparing solubilization efficiency using 1% DDM, 0.3% CHAPS, and 1% CHAPS. PMP22 was most effectively solubilized by 1% DDM. (E) Silver-stained gel of the PMP22-ALFA Co-IP workflow validating high purity of the PMP22-ALFA eluate in HEK cells. PMP22 transfected HEK cells serve as a control. 9-fold higher amounts of ALFA IP fractions were loaded as compared with input. (F) Chymotrypsin digestion confirms comparable amounts of PMP22-ALFA in the input of the experimental condition and PMP22 in the input of the control, as determined by intensity-based absolute quantification (iBAQ) intensity. Further, it validates the specific enrichment of PMP22-ALFA in the ALFA IP fraction. (G) Co-transfection of PMP22 with PMP22-GFP and PMP22-ALFA with PMP22-GFP confirms PMP22 oligomer formation via Western Blot analysis.

We then extended the LFQ-MS analysis to identify PMP22 interactors. Whereas chymotrypsin cleavage can facilitate detection of hydrophobic proteins with few trypsin cleavage sites such as PMP22 (Fischer & Poetsch, 2006), trypsin is better suited for enhanced protein coverage (Dau *et al*, 2020), and we therefore used trypsin as a protease for the remaining experiments. As expected for successful Co-IP, the resulting volcano plot was skewed, showing enrichment of multiple proteins in the PMP22-ALFA eluate compared with the untagged PMP22 background, while only few were depleted (Fig. 2A). The top enriched proteins included previously reported PMP22 interactors, among which were ER resident proteins involved in ER quality control such as Calnexin (CANX) (Dickson *et al*, 2002; Hara *et al*, 2014; Marinko *et al*, 2021) or protein disulfide-isomerase TMX1 (Luck *et al*, 2020), as well as the L-type lectin VIPL (LMAN2L) related to the previously reported interacting ERGIC-53 (LMAN1) (Marinko *et al*, 2021), both of which act as sorting receptors for ER export of transmembrane proteins. However, also proteins localized to the distal part of the secretory pathway where highly enriched, such as the lysosomal LAMP1, LAMP2 and TMEM192, the trans-Golgi recycling carrier Secretory carrier-associated membrane protein 2 (SCAMP2) as well as plasma membrane-localized Na^+^/K^+^-ATPase subunit beta-3 (ATP1B3) and Scavenger receptor class B member 1 (SCARB1). Functional annotation enrichment analysis of all potential interactors using the Gene Ontology aspect *Cellular Component* confirmed a balanced distribution across the intracellular organelles and the plasma membrane (Fig. 2B). In contrast, only a small fraction of the PPIs reported by Sanders and colleagues are with proteins with reported location at the plasma membrane (Marinko *et al*, 2021) (Suppl. Table 1). Similar to the results of Sanders and colleagues, functional annotation analysis according to *Biological Process* showed enrichment of proteins involved in processing and transport of proteins along the secretory pathway (Fig. 2B).

**Figure 2.**
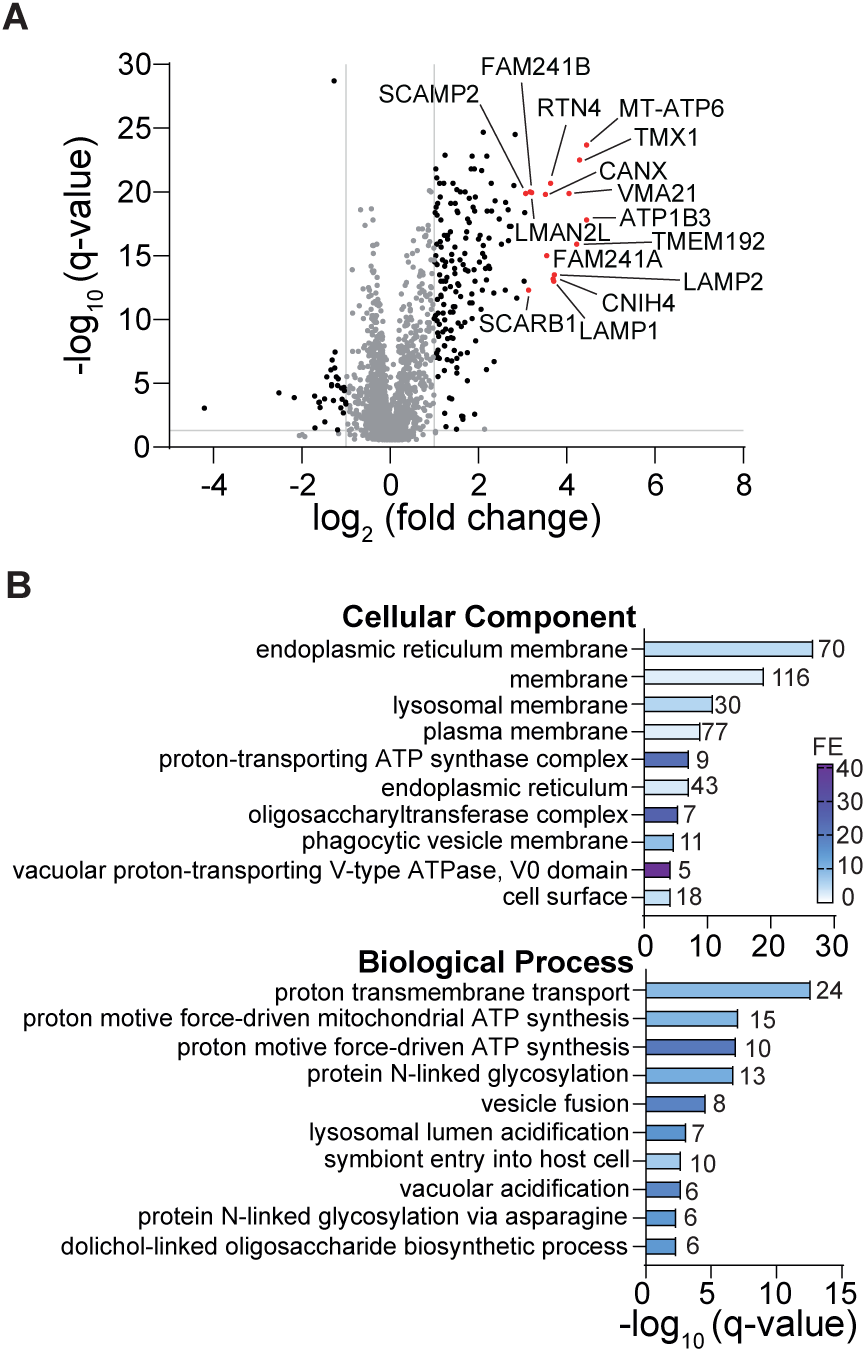
Proteomic and Go-term enrichment analysis of PMP22-ALFA-interactors in HEK cells. (A) Volcano plot of the proteins enriched in PMP22-ALFA IP over background ALFA IP with untagged PMP22. The log₂-transformed fold change is represented on the x-axis, while the −log₁₀-transformed q-value is plotted on the y-axis. The top 15 most enriched proteins in the PMP22-ALFA IP are highlighted in red. Grey lines indicate a −log₁₀-transformed q-value cutoff of 1.301 (corresponding to a q-value of 0.05) and a log₂ fold change threshold of 1, establishing the significance criteria for differential enrichment. Proteins that do not meet these thresholds are displayed in grey. (B) Functional annotation analysis of the enriched proteins for Cellular Compartment and Biological Process aspects. A q-value threshold of <0.05 was used to determine significance. The gene count and fold enrichment (FE) values are shown, with the top 10 most significantly enriched terms presented for each category. The experiment was performed on three experimental replicates and two technical replicates.

In Schwann cells, PMP22 plays a still somewhat ill-defined role as an adhesion protein in compact myelin (Pashkova *et al*, 2024; D’Urso *et al*, 1999) as well as in tight and adherens junctions of non-compact myelin (Guo *et al*, 2014; Hu *et al*, 2016). Moreover, an impact of PMP22 on integrin-mediated adhesion to the extracellular matrix (ECM) has been reported in Schwann cells (Amici *et al*, 2006; Poitelon *et al*, 2018) and other cells (Brancolini *et al*, 1999; Rao *et al*, 2011). Beyond Schwann cells in peripheral nerves, *PMP22* also shows relatively strong expression in various epithelia (Baechner *et al*, 1995; Roux *et al*, 2004) and has been shown to contribute to intercellular adhesion in the epithelial Madin-Darby Canine Kidney (MDCKII) cell line (Notterpek *et al*, 2001; Roux *et al*, 2005; Zoltewicz *et al*, 2012). Several PPIs of PMP22 related to a function in extracellular and autotypic adhesion have been reported (Amici *et al*, 2006; Guo *et al*, 2014; Hu *et al*, 2016; Pashkova *et al*, 2024). We reasoned that Co-IP-MS from MDCKII cells is suitable for discovering interaction partners of PMP22-ALFA that are related to such specialized functions as cellular adhesion. Fig. 3A shows that indeed CD47, a plasma membrane localized protein involved in integrin-mediated adhesion (Wang *et al*, 1999; Reed *et al*, 2019) and previously reported as a peripheral myelin protein (Gitik *et al*, 2011; Siems *et al*, 2020), is among the highest enriched proteins in MDCKII cells. Functional annotation analysis according to *Biological Process* further shows the term *cell adhesion* highly enriched (Fig. 3B), the most enriched PPI candidates in the PMP22-ALFA eluate of this group being CD47, the beta 1 subunit of the Na^+^/K^+^-ATPase (ATP1B1), which has a crucial function in the formation of adherens junctions (Vagin *et al*, 2006), the important tight junction component Claudin 1 (CLDN1) (Furuse *et al*, 1998) as well as CD44, which mediates cell-ECM interactions via binding various ECM components (Dzwonek & Wilczyński, 2015). According to the enrichment of PPI candidates associated with cell-ECM and intercellular adhesion, the *Cellular Component* terms *cell surface* and *basolateral plasma membrane* were enriched (Fig. 3B). In addition, the PMP22-ALFA eluate showed enrichment of several proteins involved in biosynthesis of the essential myelin lipid class of sphingolipids, reflected in enrichment of the term *ceramide biosynthetic process* in the functional annotation aspect *Biological Proces*s (Fig. 3B). Among the proteins in this group are the ceramide synthases CERS2 and CERS6, as well as all three isoforms of serine palmitoyltransferase (SPTLC1,2,3), the pacemaker enzyme in the *de novo* sphingolipid synthesis pathway (Quinville *et al*, 2021). Moreover, two isoforms of a regulatory protein (ORMDL2,3) and two further enzymes involved in sphingolipid synthesis, 3-ketodihydrosphingosine reductase (KDSR) and dihydroceramide desaturase (DEGS1), were enriched in the PMP22-ALFA eluate. Interestingly, among the top 15 most enriched proteins in the PMP22-ALFA eluate were Caveolin 1 (CAV1), which is crucial for formation of the cholesterol and sphingolipid sequestering caveolae (Örtegren *et al*, 2004; Ariotti *et al*, 2014), as well as Plasmolipin (PLLP), a tetraspan, highly abundant myelin protein in both the central and the peripheral nervous system that contains a sphingolipid binding domain and plays a role in myelin biogenesis by myelin precursor membrane formation in the secretory pathway (Bosse *et al*, 2003; Yaffe *et al*, 2015; Azzaz *et al*, 2023). Considering PMP22’s proposed role in regulation of myelin lipid metabolism (Stefanski *et al*, 2024; Silva *et al*, 2025), these novel PMP22 PPI candidates are of particular interest.

**Figure 3.**
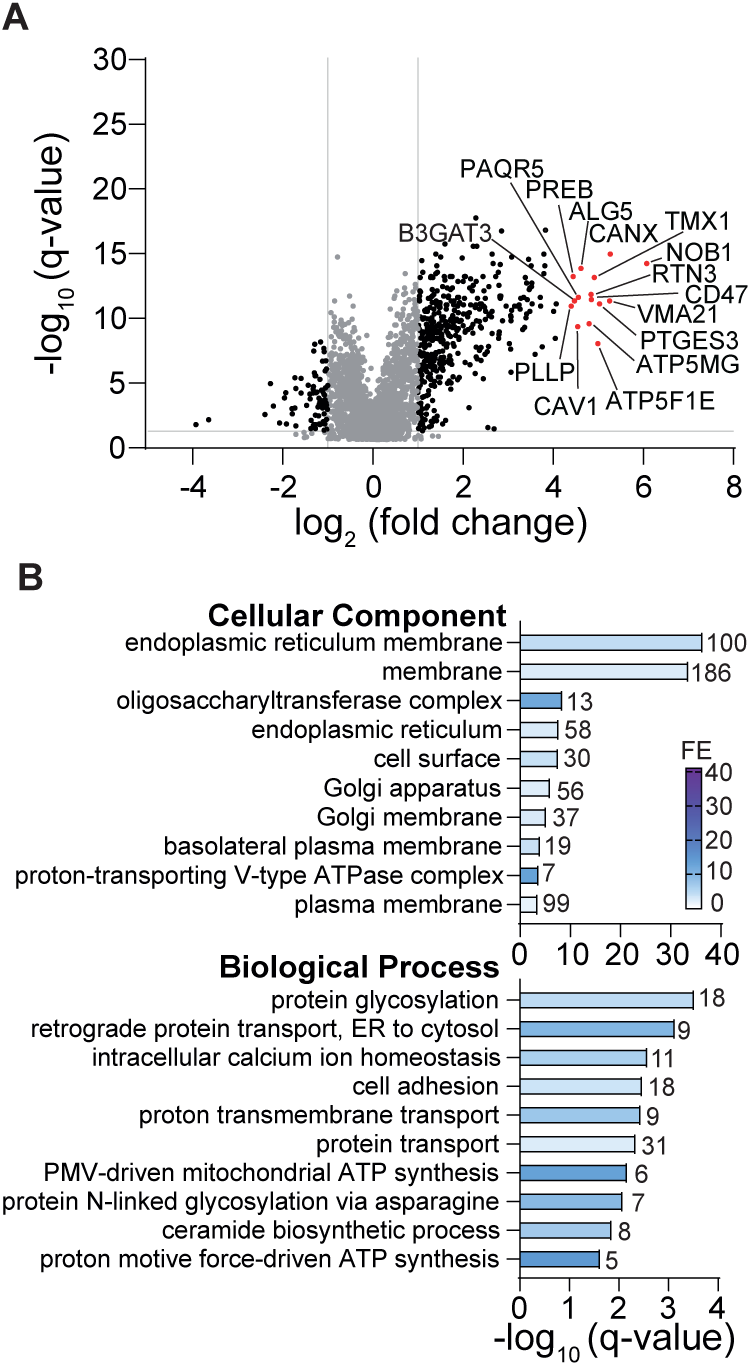
Proteomic and Go-term enrichment analysis of PMP22-ALFA-interactors in MDCKII cells. Volcano plot of the proteins enriched in PMP22-ALFA IP over background ALFA IP with untagged PMP22. The log₂-transformed fold change is represented on the x-axis, while the −log₁₀-transformed q-value is plotted on the y-axis. The top 15 most enriched proteins in the PMP22-ALFA IP are highlighted in red. Grey lines indicate a −log₁₀-transformed q-value cutoff of 1.301 (corresponding to a q-value of 0.05) and a log₂ fold change threshold of 1, establishing the significance criteria for differential enrichment. Proteins that do not meet these thresholds are displayed in grey. (B) Functional annotation analysis of the enriched proteins for Cellular Compartment and Biological Process aspects. A q-value threshold of <0.05 was used to determine significance. The gene count and fold enrichment (FE) values are shown, with the top 10 most significantly enriched terms presented for each category. The experiment was performed on three experimental replicates and two technical replicates.

PMP22 plays its most significant role in myelinating Schwann cells, reflected in the striking PMP22 dose sensitivity leading to both CMT1A and HNPP. To discover new PPI candidates in Schwann cells, we turned to MSC80, an immortalized cell line derived from secondary, peripheral nerve cell culture that retained important characteristics like Schwann cell morphology and expression of Schwann cell markers (Boutry *et al*, 1992). Importantly, MSC80 cells were able to myelinate injured axons after transplantation *in vivo* (Boutry *et al*, 1992). We treated the cells with cyclic AMP analog dbcAMP to stimulate upregulated expression of myelination related genes (Brockes *et al*, 1979; Arthur-Farraj *et al*, 2011). However, after 24 hours of dbcAMP treatment, at the proteomic level we were unable to detect any switch to a pro-myelinating expression signature (suppl. Fig. 2), which may reflect a limitation of this cell line as a Schwann cell model. Nonetheless, in stimulated MSC80 cells, several of the top 15 PPIs (Fig. 4A) candidates have previously been reported as peripheral myelin proteins (Siems *et al*, 2020), including, interestingly, those primarily associated with cellular energy regeneration in mitochondria (ATP5F1B, ATP5F1D, ATP5MF) or protein glycosylation in the ER (DDOST, RPN1). However, many canonical proteins of the myelin sheath were also found among the enriched proteins in the PMP22-ALFA eluate, including the established PMP22 interaction partner MPZ (Pashkova *et al*, 2024), the proteolipid protein (PLP1), the Rho-type GTPase cell division cycle 42 (CDC42), Cell surface glycoprotein MUC18 (MCAM) as well as several Annexin isoforms (ANXA1,2,5,6) (Shih *et al*, 1998; Benninger *et al*, 2007; Hayashi *et al*, 2007; Siems *et al*, 2020) (Fig. 4A). In line with these observations, functional annotation analysis showed enrichment of the *Cellular Component* term *myelin sheath* (Fig. 4B). Moreover, like in MDCKII cells (Fig. 3) proteins involved in sphingolipid synthesis such as SPTLC1, SPTLC2, CERS2 and KDSR were again enriched among the PMP22 PPI candidates (Fig. 4B).

**Figure 4.**
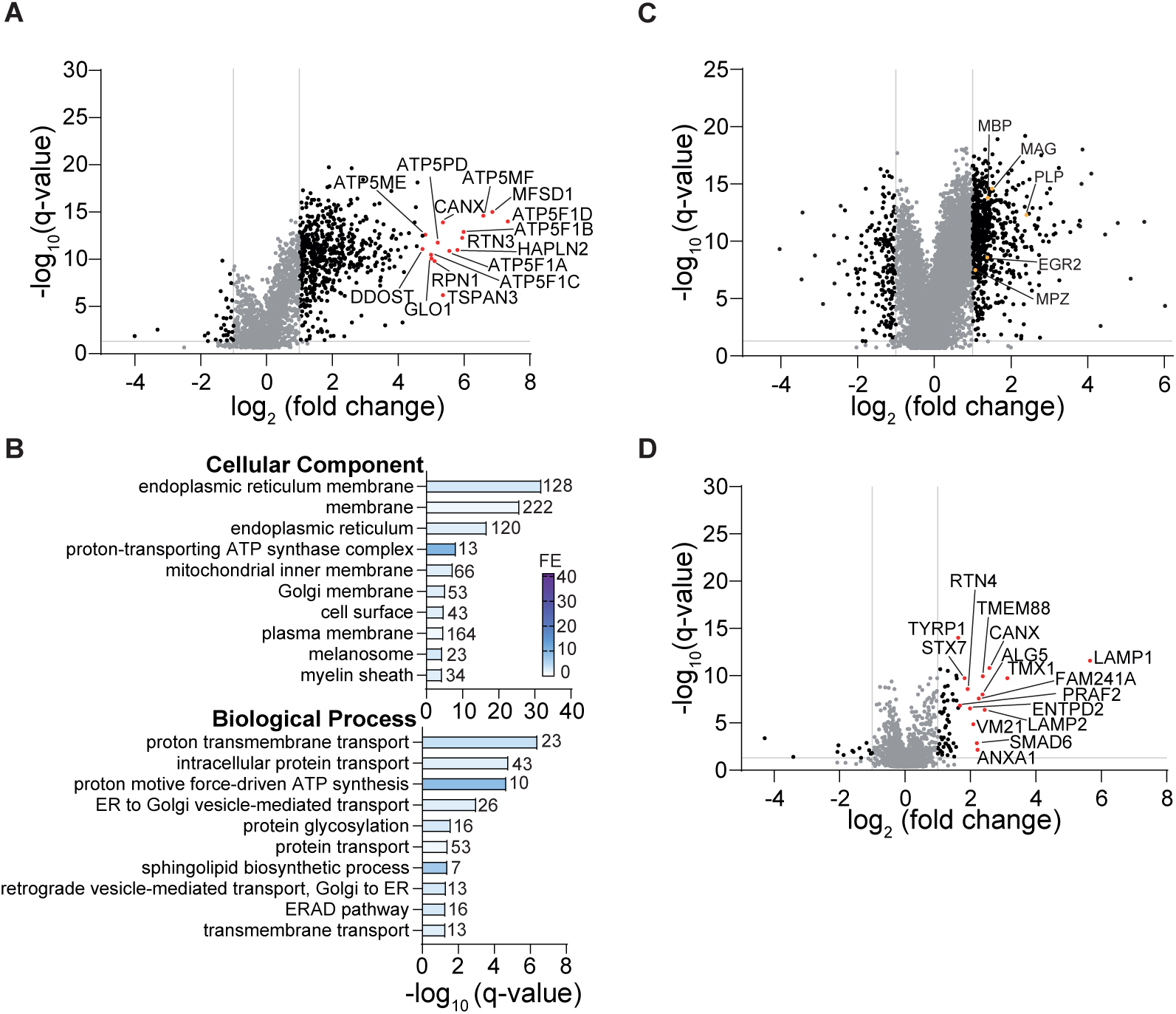
Proteomic and Go-term enrichment analysis of PMP22-ALFA interactors in Schwann cells. (A) Volcano plot of proteins identified in PMP22-ALFA IP from MSC80 cells, compared to control IP with untagged PMP22. The top 15 enriched proteins are highlighted in red. (B) Functional annotation analysis of proteins enriched in the PMP22-ALFA IP from MSC80 cells, covering the categories Cellular Compartment and Biological Process. The top 10 significantly enriched terms (q < 0.1) are shown with gene counts and fold enrichment (FE) values. (C) Differential proteome analysis of primary Schwann cells under unstimulated and cAMP-stimulated conditions. Key myelin proteins (MBP, MAG, PLP, EGR2, MPZ) are highlighted in orange. (D) Volcano plots of proteins identified in PMP22-ALFA IP from stimulated primary Schwann cells, compared to control IP with untagged PMP22. Volcano plots display log₂ fold change on the x-axis versus –log₁₀ q-value on the y-axis. Grey lines indicate the thresholds for significant enrichment (q < 0.05; log₂ fold change > 1). Proteins not meeting these criteria are shown in grey. All experiments were performed in three biological and two technical replicates.

We then turned to primary Schwann cells isolated from peripheral nerves of rats at postnatal day (P) 2-P4 since we expected primary Schwann cells to be more readily stimulated towards a differentiated phenotype than the MSC80 cell line. Indeed, upon dbcAMP treatment, we found the pro-myelinating transcription factor Krox20 (EGR2) together with major myelin proteins upregulated on the protein level (Fig. 4C). However, as compared with the previously investigated cell types, transient transfection was only successful in a smaller fraction of primary Schwann cells (suppl. Fig. 1D, D’). Thus, probably due to high background from non-transfected cells contributing to the Co-IP input, we found considerably less (63) proteins that were enriched in the PMP22-ALFA eluate (Fig. 4D, Fig. 5A), and functional annotation did not show any significant enrichment. Despite the considerably lower number of PPI candidates that resulted from primary Schwann cells as compared to other cell types, we again found multiple reported myelin proteins including those that we found in MSC80 cells, such as MPZ, PLP1, MCAM and ANXA1. The Venn diagram in Fig. 5A shows the overlap of all PPI candidates shown in Figs. 2-4, and Fig. 5B lists all PPI candidates that were found at least in two cell types.

**Figure 5.**
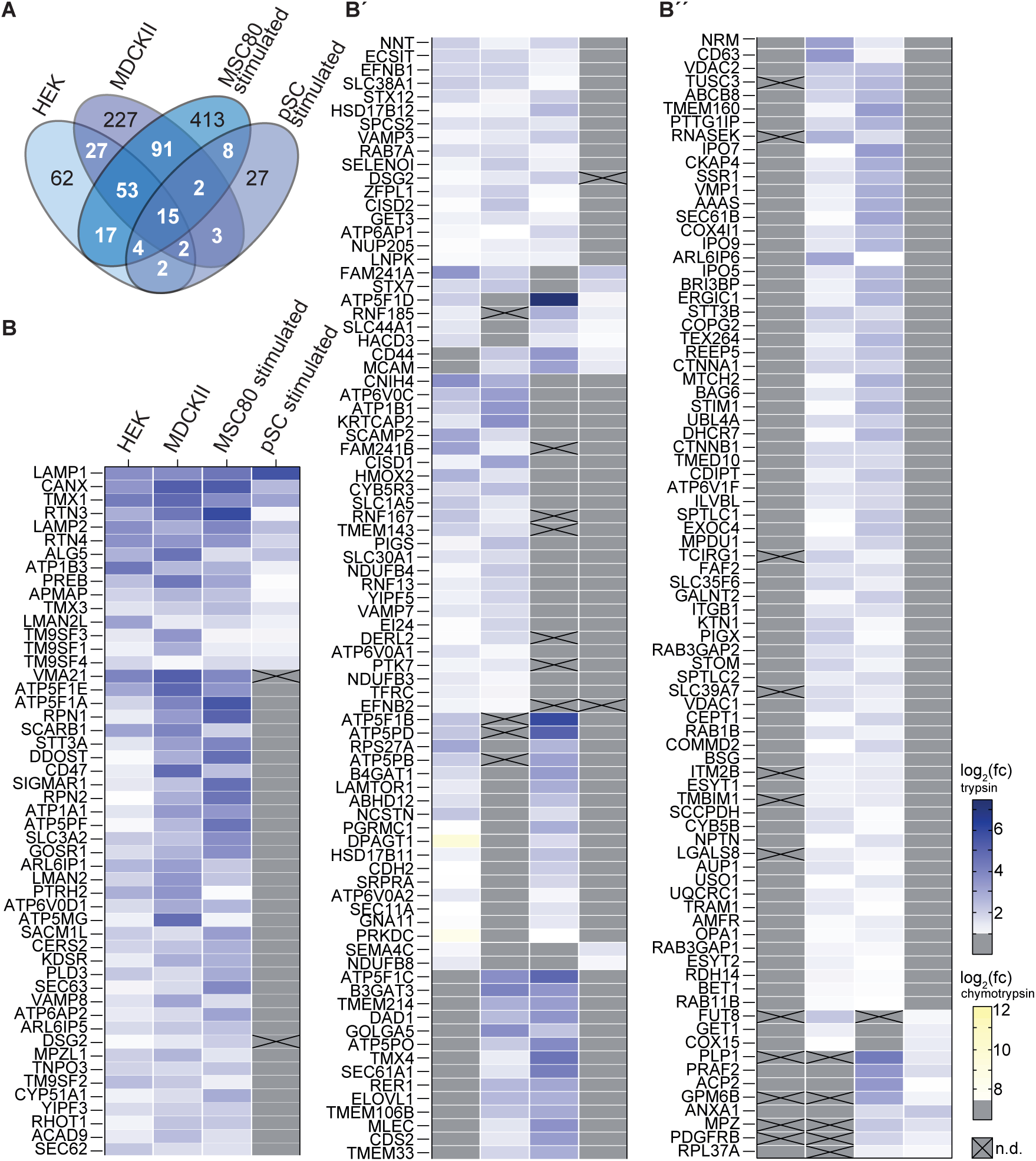
Comparison of PMP22-ALFA PPIs across cell types and conditions. (A) Venn diagram illustrating the overlap of identified proteins across four PMP22 ALFA Co-IP conditions, considering proteins with a q-value below 0.05 and a log₂-transformed fold change >1. The four groups analyzed are HEK cells, MDCKII cells, stimulated MSC80 and primary Schwann cells (pSC). (B), (B’) and (B’’) List of proteins identified in the Co-IP experiments including those detected in at least two of the cell types/conditions, with a q-value below 0.05. The heat map represents the log₂ fold change of each protein relative to the control, with values indicated by shades of blue (trypsin digestion) or yellow (chymotrypsin digestion, shown for proteins that were not detected after trypsin digestion). Grey indicates that the protein was not detected or did not meet the cutoff criteria of enrichment in the ALFA-IP, while a cross denotes that it was also not detected in the respective input sample.

## Discussion

The aim of this study was to identify novel PPIs of PMP22, with a view to providing a basis for further, functional investigations. We employed a Co-IP approach combining the optimized, efficient and gentle removal of PMP22 from its native environment in various cellular membrane compartments including the plasma membrane, with a highly specific, high-affinity epitope-nanobody pair. This allowed 1,074 new candidates to be added and confirming 54 of the existing—to our knowledge—296 PMP22 candidate PPIs (suppl. Table 2). In the extensive data available on the interactome of eukaryotic cells derived from the most successful techniques for large-scale PPI discovery, Y2H and AP-MS/IP-MS, PPIs of membrane proteins are underrepresented, due to various biological and technical challenges such as misfolding and subcellular localization as well as insufficient solubilization (Sharifi Tabar *et al*, 2022). It is therefore not surprising that the PPIs of PMP22 that have been discovered using these techniques to date are relatively few compared to our results (Wang *et al*, 2011, 2023; Dittmer *et al*, 2014; Rolland *et al*, 2014; Sahni *et al*, 2015; Luck *et al*, 2020; Marinko *et al*, 2021; Haenig *et al*, 2020). Through efficient solubilization of PMP22, most likely this study was the first to enable sampling of its PPIs in all cellular membrane compartments. Thus, besides intracellular compartments like ER and Golgi, in functional annotation analysis we found a significant fraction of PPI candidates localized at the cell surface (Figs. 2B, 3B, 4B), including specialized plasma membrane compartments like the basolateral membrane of MDCKII cells where intercellular junctions are formed (Fig. 3B). In the Schwann cell line MSC80 we found the term myelin sheath enriched (Fig. 4B), with several known myelin proteins like PLP1, MPZ, MCAM and ANXA1 enriched in the PMP22-ALFA eluates of MSC80 and primary Schwann cells. At the onset of myelination, Schwann cells mount a transcriptional program enabling coordinated synthesis of both myelin proteins and lipids that are required in large quantities for myelin sheath formation (LeBlanc *et al*, 2005; Pertusa *et al*, 2007; Fledrich *et al*, 2018; Kim *et al*, 2018; Poitelon *et al*, 2020). In recent years, evidence has accumulated that links PMP22 to lipid metabolism and importantly, dysregulation of lipid synthesis and transport to PMP22-related neuropathies, most severely affecting the major myelin sterol lipid cholesterol as well as the lipid class of sphingolipids (Fledrich *et al*, 2012, 2018; Zhou *et al*, 2019; Visigalli *et al*, 2020; Michailidou *et al*, 2023; Prior *et al*, 2024; Capodivento *et al*, 2024; Zhou *et al*, 2020). Strikingly, both in polarized epithelial cells and in Schwann cells, we here detected interaction of PMP22 with the sphingolipid synthesis machinery of the ER. The initial step of the *de novo* sphingolipid synthesis pathway is catalyzed by the serine palmitoyltransferase complex (SPT), a homo-dimeric assembly of a hetero-tetramer that consists of the enzyme core (SPTLC1 and SPTLC2 or SPTLC3) and two transmembrane accessory subunits (ORMDL1,2 or 3 and serine palmitoyltransferase small subunit (SPTSS) A or B) (Li *et al*, 2021; Wang *et al*, 2021). All SPT subunits resulted as PMP22 PPI candidates but SPTSSA and SPTSSB, which are only 71 or 76 amino acids in length and feature unfavorable tryptic cleavage sites for detection via MS. Accordingly, we could not detect the small subunits in any of the Co-IP input samples. Of note, Co-IP from cell lysates results in enrichment of both direct and indirect interactors, and interaction with just one of the subunits of a stable complex can thus suffice to detect all subunits. To our knowledge, there is no evidence so far that the remaining enzymes of the *de novo* sphingolipid synthesis pathway (CERS, KDSR, DEGS) interact with SPT, so they may operate separately. Since in addition to SPT, we identified all three as PPI candidates, it seems likely that PMP22 can directly interfere with sphingolipid synthesis in the ER, with a possible functional role of contributing to physiological regulation of sphingolipid supply to the myelin sheath and to its dysregulation in CMT1A. PPIs resulting from Co-IP-MS inevitably contain false positives due to contaminants or background noise (Dunham *et al*, 2012). Therefore, our PPI results cannot be considered valid on their own. Instead, when used as input for future candidate or functional screening studies, PPI candidates will have to be validated using orthogonal technical approaches. These might include techniques providing evidence of physical interaction like crosslinking MS (Larance *et al*, 2016; Leitner *et al*, 2016; O’Reilly & Rappsilber, 2018; Piersimoni *et al*, 2022) or more direct methods that require purified proteins like surface plasmon resonance (Puiu & Bala, 2016). Proximity of PMP22 and PPI candidates *in-situ* may be tested using techniques that also provide information about the subcellular location of PPI, like Proximity ligation in fixed cells (Söderberg *et al*, 2006) or FRET/BRET assays in living cells (Machleidt *et al*, 2015). In light of the dosage effect of PMP22, we also note that transient overexpression can be expected to result in considerable alterations in its PPIs. Moreover, by restricting our approach to monoculture *in vitro*, we certainly have missed those interactions of PMP22 that require the cellular architecture of myelinating Schwann cells *in vivo*. Future investigations employing DNA-editing approaches may be able to sample the entirety of PMP22’s PPIs, at endogenous expression levels within the developing axon-Schwann cell unit. Nevertheless, in this study we were able to significantly expand the range of potential interaction partners of PMP22, thereby creating new, evidence-based starting points for future studies. Given that there is still no therapy available in CMT1A, the identification of new PPIs may thus form a new family of future therapeutic targets for PMP22-associated diseases.

## Methods

### Cell culture

HEK293T cells (MERCK, #96121229) and MSC80 cells (Boutry et al., 1992) were cultured in DMEM (Gibco, Thermo Fisher) supplemented with 10% FCS (Gibco, Thermo Fisher) and 1% penicillin/streptomycin (Pen/Strep (Lonza)) while MDCKII cells (Merck, #00062107) were maintained in MEM medium (Gibco, Thermo Fisher), supplemented with 5% FCS and 1% Pen/Strep.

Primary Schwann cell monocultures were prepared from rats at P2-P4. Dissected sciatic nerves of six rats were pooled and enzymatically digested with trypsin (Invitrogen, Thermo Fisher) and collagenase II (Worthington) for 1 hour at 37 °C. The tissue was dissociated by triturating and the reaction stopped by adding 25% FCS in basic medium (DMEM). Cells were plated in Schwann cell basic medium (1% Pen/Strep, 10% FCS, 1% GlutaMax (Gibco, Thermo Fisher)) on PLL-coated culture plates. The following day fibroblasts were eliminated by supplementing the medium with 10 µM Cytosine β-D-arabinofuranoside hydrochloride (AraC (Sigma)) for 2 days. Primary Schwann cells were stimulated for 24 hours with 2 mM dbcAMP (BIOLOG) in the presence of 5% FCS, control cells were maintained in growth medium containing 5% FCS for this incubation period.

HEK293T cells were transiently transfected with ALFA-, GFP-, or untagged PMP22 expression plasmids (VectorBuilder) using polyethylenimine (Polysciences, Inc), while MDCKII, MSC80, and primary Schwann cells were transfected via Lipofectamine™ 3000 (Thermo Fisher).

### Cell lysates

Cell pellets from one transfected 10 cm plate were resuspended in ice-cold TBS buffer (100 mM Tris/HCl, 150 mM NaCl, pH 7.5) supplemented with a complete protease inhibitor cocktail (Roche Diagnostics). To compare optimal detergents for analysis, samples were incubated for 1 hour at RT with either 1% CHAPS (AppliChem), 0.3% CHAPS, or 1% DDM (Anatrace), followed by centrifugation at 100,000 × g for 1 hour at 4 °C.

### Co-IP

DDM-solubilized cell lysate was diluted with TBS to a final concentration of 0.15% DDM. Insoluble material was pelleted via centrifugation at 100,000 × g for 1 hour at 4 °C. Protein concentrations were determined with the Bio-Rad DC™ Protein Assay Kit and adjusted. Magnetic ALFA Selector PE Resin beads were washed and equilibrated before adding the cleared cell lysate. After incubation for 90 minutes at 4 °C, the beads were washed six times with cold TBS buffer containing 0.025% DDM. PMP22-ALFA was then eluted by providing an excess of ALFA peptide.

### Glycosylation assay

Cleared cell lysates were treated with Endoglycosidase (Endo) H and Peptide N-Glycosidase (PNGase) F (New England Biolabs) according to the manufacturer’s instructions, and the proteins were evaluated by Western blotting with peroxidase-conjugated anti-ALFA antibody.

### SDS Page and Western Blotting

Proteins were separated by electrophoresis using 4–20% precast polyacrylamide gels (Novex™ Tris-Glycin Mini gels, Thermo Fisher). PageRuler Plus Prestained Protein Ladder (Thermo Fisher) was used for loading and size control. The proteins were then transferred to a methanol-activated PVDF membrane (Immobilon-P Membrane; Millipore, Merck). Fast Green staining was performed to determine the total protein load. Therefore, membranes were rinsed in Fast Green washing solution (30% methanol, 6.7% glacial acetic acid) to remove transfer buffer residues and then incubated in Fast Green staining solution (0.5% Fast Green (Sigma), 30% methanol, 6.7% glacial acetic acid) for 5 minutes. The membranes were then washed twice in Fast Green washing solution before fluorescent imaging using the ChemoStar imager (INTAS Science Imaging Instruments). Membranes were blocked in milk-TBST (5% milk powder, 25 mM Tris, 75 mM NaCl, 0,05 % Tween 20) for 1 hour at room temperature, and then incubated overnight at 4 °C with primary antibodies (sdAB ALFA-HRP 1:40000; NanoTag Biotechnologies N1501 #15201101; GFP 1:1000, Abcam #ab290), diluted in blocking solution. After 5–7 washing steps with TBST, membranes were incubated, when required, with HRP-conjugated secondary antibodies (1:20000, Dianova), for 2 hours at room temperature, followed by additional washes and detection using the Western Lightning Plus ECL Kit.

### Silver staining

For silver staining, gels were fixed overnight in 40% ethanol and 10% acetic acid. The next day the gels were washed twice for 20 minutes in 20% ethanol, and once for 20 minutes in ddH₂O. Pretreatment was performed for 1 minute in 0.8 mM sodium thiosulfate, followed by three 20 seconds washes in ddH₂O. Gels were then stained for 20 minutes in 0.2% silver nitrate containing 0.02% formaldehyde and washed again three times for 20 seconds in ddH₂O. Protein bands were developed in 3% sodium carbonate with 0.02% formaldehyde until clearly visible, and the reaction was stopped by two 10 minutes incubations in 5% acetic acid.

### Immunofluorescence

PMP22-ALFA transfected HEK293T cells were fixed in 4% PFA, washed three times with PBS, and permeabilized in ice-cold methanol/acetone (95% / 5%) for 5 minutes, following another washing series of three times with PBS. Coverslips were placed on blocking solution (2% horse serum (Gibco, Thermo Fisher), 2% BSA (BioMol), 0.1% gelatine) and incubated for 1 hour. Coverslips were incubated overnight at 4 °C with the primary antibody (ALFA polyclonal, 1:500; NanoTag Biotechnologies, N1581 #082312) diluted in blocking solution. Following this, the coverslips were washed three times with PBS and incubated with secondary antibodies (1:1000) for 1 hour at room temperature. After another set of washes, coverslips were mounted onto microscopic slides and imaged using the Zeiss Axio Imager Z1 microscope.

### Sample preparation for proteomic analysis

HEK293T, MSC80, and MDCKII cells, as well as primary rat Schwann cells were lysed and in-solution digested by trypsin followed by the data independent acquisition (DIA) proteomic pipeline.

Samples of lysed cells or eluates from ALFA beads were digested by trypsin and cleaned up using SP3 paramagnetic bead protocol (Hughes *et al*, 2019). Briefly, 50 μL of lysates or eluates containing approximately 15 μg total protein was mixed 1:1 (v/v) with 50 mM ammonia bicarbonate and incubated for 10 min at 37°C and 900 rpm shaking. The disulfide bonds were reduced with 10 mM DTT for 30min at 37°*C* and 900 rpm followed by alkylation with 40 mM iodoacetamide (IAA) for 30 min at 25°C and 900 rpm and quenching of the alkylating agent with 10 mM DTT for additional 10 min at *25°*C and 900 rpm. The SP3 beads (PreOmics) were prepared and the samples were bound to the beads as described before using a magnetic rack for Eppendorf tubes. Samples were pre-digested by Benzonase (Universal Nuclease for Cell Lysis, 100kU, Pierce) for 1h at 37°C and 1000 rpm and then by trypsin (Sequencing Grade Trypsin, Promega) in 1:50 w/w ratio to protein overnight at 37°C and 1000 rpm. Tryptic peptides were eluted from the beads according to the SP3 protocol and the resultant eluates were dried in the vacuum concentrator (Eppendorf). Dried samples were redissolved in 0.1% (v/v) formic acid and 2% (v/v) acetonitrile solution. For each run of LC-MS/MS analysis, 100 ng of total protein was injected as measured in lysates before trypsin digestion.

For DIA pipeline, samples were analyzed in 3 technical replicates in each group of samples (PMP22 transfected and control input cells and corresponding IP eluates) and in 2 measurement replicates for each technical replicate.

Since the protein of interest, PMP22, has a low amount of trypsin digestion sites which leads to formation of few long peptides with low detection levels in LC-MS/MS, HEK293T sample set was digested by chymotrypsin followed by the data dependent acquisition (DDA) proteomic pipeline. Input and IP eluate samples were separated by conventional SDS-PAGE in triplicates. Each line was cut into equal 23 slices and the slices were in-gel digested by chymotrypsin (sequencing grade, Promega) as described earlier (Shevchenko *et al*, 1996).

### LC-MS/MS analysis (DIA pipeline)

Nano-HPLC was performed using Dionex UltiMate 3000 chromatographer, where LC effluent solutions were Buffer A (0.1% (v) formic acid) and Buffer B (80% (v) ACN, 0.08% (v) formic acid). The columns used in LC were the Trap column (PEPMAP100 C18, 0.3 x 5 mm, 5 µm, Thermo) and the main column (Aurora CSI C18, 25 cm x 75 µm, 1.6 µm; IonOpticks). HPLC was performed under a column temperature of 50°C. The solution gradient was as follows: 0-3.5 min 0.4 µL/min 5% (v) buffer B, 3.5-5 min 0.4-0.2 µL/min 5-10% B, 5-40 min 0.2 µL/min 10-42% B, 40-41 min 0.2 µL/min 42-95% B, 41-45 min 0.2 µL/min 95% B, 45-45.5 min 0.2-0.4 µL/min 95-5% B, 45.5-50 min 0.4 µL/min 5% B, where the flow and % buffer B were changing linearly.

The chromatographer was coupled with timsTOF Pro 2 mass spectrometer (Bruker Daltonics). The applied ionization method was nanoelectrospray (Bruker CaptiveSpray) with positive ionization polarity where the capillary voltage was 1.5 kV and dry gas flow rate was 3 L/min. Ionization took place at 180 °C. The TIMS chamber was filled with nitrogen gas at 305 K temperature. The pressure at the tunnel entrance was ∼2.7 mbar.

For data-independent acquisition in dia-PASEF high-sensitivity mode (the default method provided by the vendor), an ion mobility range was sampled from 1/K0 1.43 to 0.6 Vs/cm2 using equal ion accumulation and ramp times in the dual TIMS analyzer of 100 ms each. The collision energy was lowered as a function of increasing ion mobility from 59 eV at 1/K0 1.43 Vs/cm2 to 20 eV at 1/K0 0.6 Vs/cm2. Overall, 16 PASEF scans were used with two mass windows per the scan.

The MS and TIMS were calibrated linearly using three ions from the Agilent ESI LC/MS tuning mix (m/z, 1/K0: 622.0289 Th, 0.9915 V·s·cm^−2^; 922.0097 Th, 1.1986 V·s·cm^−2^; 1221.9906 Th, 1.3934 V·s·cm^−2^) in positive mode.

### LC-MS/MS analysis (DDA pipeline)

Peptides after in-gel digestion of chymotrypsin were loaded onto nano-HPLC (Dionex Ultimate 3000 UHPLC Thermo Fischer Scientific) coupled with in-house packed C18 column (ReproSil-Pur 120 C18-AQ, 3 µm particle size, 75 µm inner diameter, 30 cm length, Dr. Maisch GmbH). The peptides were separated with a linear gradient of 11–40% buffer B (80% acetonitrile and 0.1% formic acid) at flow rate of 300 nL/min over 37 min gradient time at overall method duration of 58 min. Eluting peptides were analysed by Orbitrap Exploris 480 mass spectrometer (Thermo Fischer Scientific). The following MS settings were used: MS1 scan range, 350–1400 m/z; MS1 resolution, 60,000 FWHM; AGC target MS1, custom; maximum injection time MS1, custom; intensity threshold, 1E4; isolation window, 1.6 Th; normalized collision energy, 28%; charge states, 2+ to 6+; dynamic exclusion, custom; top 20 most abundant precursors were selected for fragmentation; MS2 resolution, 15,000, AGC target MS2, custom; maximum injection time MS2, custom.

### LC-MS/MS data processing (DIA pipeline)

Quantitative analysis of proteins from DIA LC-MS/MS runs was done using Spectronaut software (version 19.6.250122.62635, Biognosys) in directDIA mode with default parameters except data imputation which was on with a “background signal” option. Search databases represented reference proteome fasta files downloaded from Uniprot (Bateman *et al*, 2025): human (UP000005640, 2024-02-20), dog (UP000805418, 2024-12-04), rat (UP000002494, 2025-01-22), mouse (UP000000589, 2025-07-16).

### LC-MS/MS data processing (DDA pipeline)

Data were processed by MaxQuant (v. 1.6.17.0) (Cox & Mann, 2008) with default parameters for DDA except chymotrypsin as a protease. iBAQ values were used for protein quantitation. Differentially expressed proteins were identified using DEP2 R package (Feng *et al*, 2023).

### PPI and functional annotation enrichment analysis

The following criteria were utilized to determine potential PMP22-ALFA interacting proteins. A false discovery rate (FDR)-corrected q-value of less than 0.05 was applied to define significant changes in protein abundance. Only proteins that showed a >1 log2 fold increase in abundance between PMP22-ALFA and control IP samples were considered. To exclude background effects, proteins with a log2 fold change >0.58 (1.5-fold) in the corresponding input samples were removed. Functional enrichment of proteins was performed using DAVID (Database for Annotation, Visualization, and Integrated Discovery)(Sherman *et al*, 2022; Huang *et al*, 2009). UniProt IDs of PMP22 PPI candidates were submitted to DAVID, with input proteins serving as the background. To ensure specificity, enrichment and analysis was carried out utilizing the GO Direct category.

## Data availability

The mass-spectrometry data generated for this study are deposited in the MassIVE repository and submitted to ProteomeXchange Consortium (Deutsch *et al*, 2023) and are available using the following identifiers MSV000099338 (MassIVE), PXD069006 (ProteomeXchange).

## Acknowledgments

H.U. was funded by the Deutsche Forschungsgemeinschaft (DFG) SFB1565 (project P04, project number 469281184). K.A.N. was funded by the DFG (NA 262/3-1). M.W.S. was funded by the DFG (SE 1944/3-1). The authors thank Annika Reinelt (Bioanalytical Mass Spectrometry, Max Planck Institute for Multidisciplinary Sciences), Michael Kothe (Department of Neurogenetics, Max Planck Institute for Multidisciplinary Sciences) and Beate Veith (Department of Neurology, University Medical Center Göttingen) for their excellent technical assistance.

## Notes

### Competing Interest Statement

The authors have declared no competing interest.

### Summary of Updates

Author affiliations updated. No additional changes made

