## Supplementary Figures for "High affinity cross-context cellular assays reveal novel protein-protein interactions of peripheral myelin protein of 22 kDa"

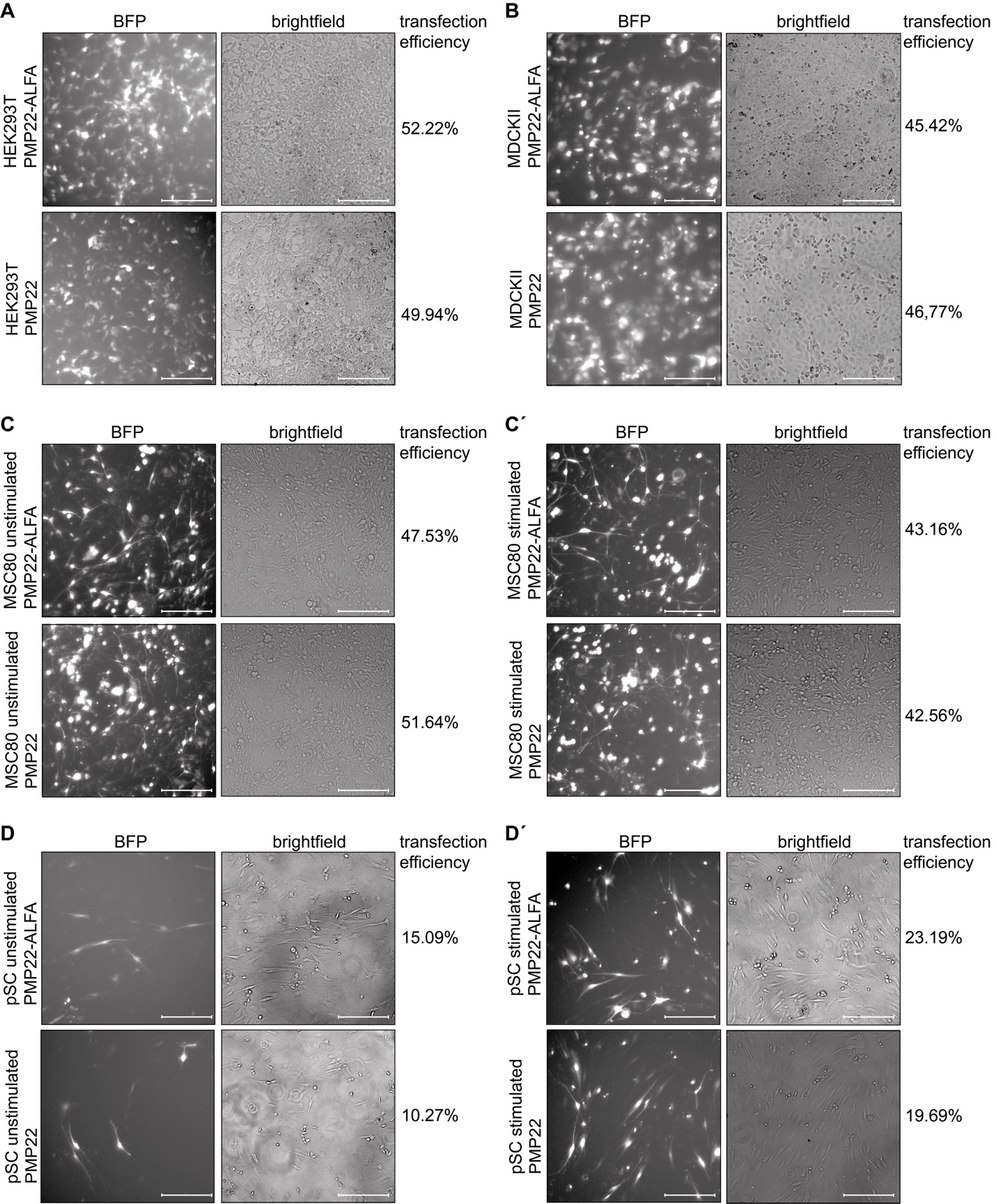
**Supplementary Figure 1. Transfection efficiency of PMP22 and PMP22-ALFA constructs across different cell types.** Representative images showing transfection efficiency of PMP22 and PMP22-ALFA in (A) HEK293T cells, (B) MDCKII cells, MSC80 Schwann cells under (C) unstimulated and (C′) cAMP-stimulated conditions, and primary Schwann cells (pSCs) under (D) unstimulated and (D′) cAMP-stimulated conditions. BFP fluorescence (left) indicates successfully transfected cells, with corresponding brightfield images shown on the right. The percentage of BFP-positive cells is indicated for each condition.

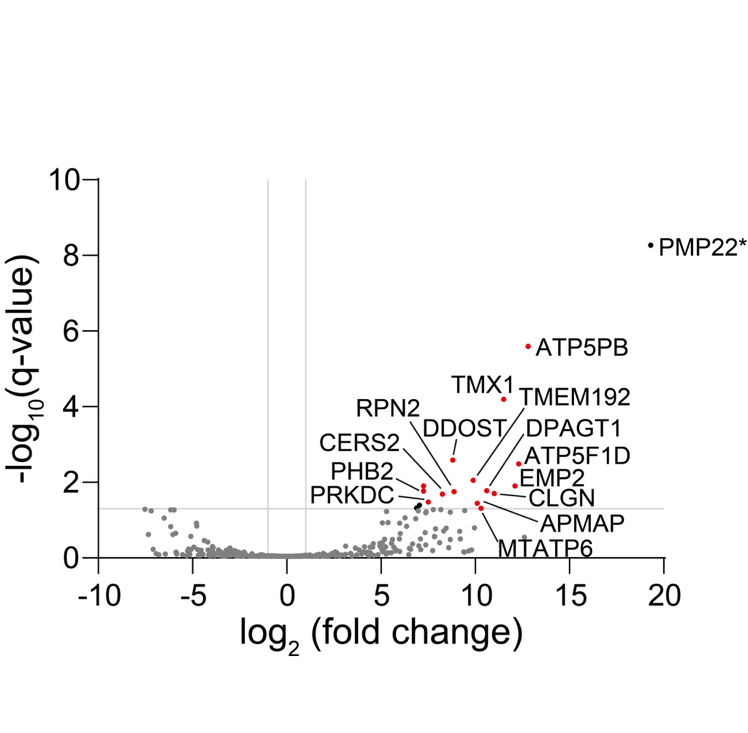

**Supplementary Figure 2. Proteins enriched in PMP22-ALFA IP from HEK cells following chymotrypsin digestion.** Volcano plot illustrating proteins enriched in PMP22-ALFA IP relative to background ALFA-IP with untagged PMP22. The log₂-transformed fold change is represented on the x-axis, while the -log₁₀-transformed q-value is plotted on the y-axis. The 15 most enriched proteins in the PMP22-ALFA IP are highlighted in red. The top 15 most enriched proteins in the PMP22-ALFA IP are highlighted in red. Grey lines indicate a -log₁₀-transformed q-value cutoff of 1.301 (corresponding to a q-value of 0.05) and a log₂ fold change threshold of 1. Proteins that do not meet these thresholds are displayed in grey. *PMP22 enrichment reflects both endogenously expressed PMP22 and recovery of transiently overexpressed PMP22-ALFA bait.

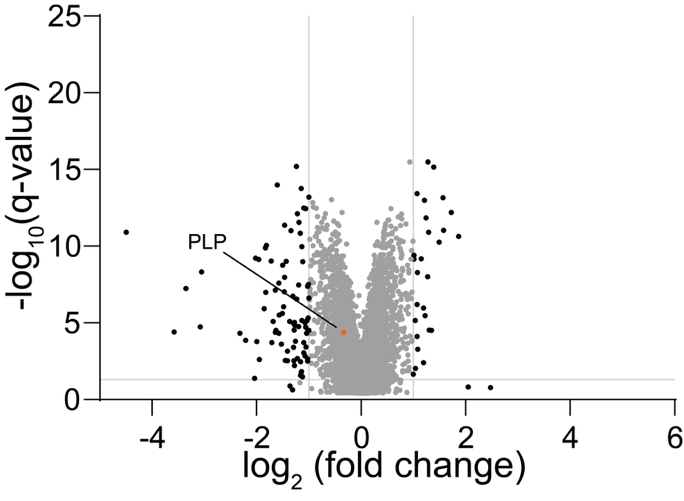

**Supplementary Figure 3. Limited proteomic response of MSC80 Schwann cells to dbcAMP treatment.** Differential proteome analysis of MSC80 Schwann cells under cAMP-stimulated vs. unstimulated conditions. The x-axis of the volcano plot displays the log₂-transformed fold change, while the y-axis represents the –log₁₀-transformed q-value. Grey lines indicate the thresholds for significant enrichment (q < 0.05; log₂ fold change > 1). Proteins not meeting these criteria are shown in grey. All experiments were performed in three biological and two technical replicates. PLP is highlighted in orange as the only protein detected in this dataset that was highlighted as significantly enriched after cAMP stimulation in primary Schwann cells (Fig. 4C).
